# Inactivation of the monofunctional peptidoglycan glycosyltransferase SgtB allows *Staphylococcus aureus* to survive in the absence of lipoteichoic acid

**DOI:** 10.1101/311282

**Authors:** Eleni Karinou, Christopher F. Schuster, Manuel Pazos, Waldemar Vollmer, Angelika Gründling

## Abstract

The cell wall of *Staphylococcus aureus* is composed of peptidoglycan and the anionic polymers lipoteichoic acid (LTA) and wall teichoic acid. LTA is required for growth and normal cell morphology in *S. aureus.* Strains lacking LTA are usually only viable when grown under osmotically stabilizing conditions or after the acquisition of compensatory mutations. LTA negative suppressor strains with inactivating mutations in *gdpP*, resulting in an increase in intracellular c-di-AMP levels, have been described previously. Here, we sought to identify factors other than c-di-AMP that allow *S. aureus* to survive without LTA. LTA-negative strains able to grow in un-supplemented medium were obtained and found to contain mutations in *sgtB, mazE, clpX* or *vraT*. The growth improvement through mutations in *mazE* and *sgtB* was confirmed by complementation analysis. We also show that an *S. aureus sgtB* transposon mutant, inactivated for the monofunctional peptidoglycan glycosyltransferase SgtB, displays a 4-fold increase in the MIC towards a number of cell wall-targeting antibiotics, suggesting that alteration in the peptidoglycan structure could help bacteria compensate for the lack of LTA. Muropeptide analysis of peptidoglycan isolated from a WT and *sgtB* mutant strains did not reveal any sizable alternations in the peptidoglycan structure. In contrast, the peptidoglycan isolated from an LTA-negative *ltaS* mutant strain showed a significant reduction in the fraction of highly crosslinked peptidoglycan, which was partially rescued in the *sgtB*/*ltaS* double mutant suppressor strain. Taken together, these data point towards an important function of LTA in cell wall integrity through its requirement for proper peptidoglycan assembly.

**Importance:** The bacterial cell wall acts as primary defence against environmental insults such as changes in osmolarity. It is also a vulnerable structure as defects in its synthesis can lead to growth arrest or cell death. The important human pathogen *Staphylococcus aureus* has a typical Gram-positive cell wall, which consists of peptidoglycan and the anionic polymers lipoteichoic acid (LTA) and wall teichoic acid. Several clinically relevant antibiotics inhibit the synthesis of peptidoglycan; hence it and teichoic acids are considered attractive targets for the development of new antimicrobials. We show that LTA is required for efficient peptidoglycan crosslinking in *S. aureus* and inactivation of a peptidoglycan glycosyltransferase can partially rescue this defect, altogether revealing an intimate link between peptidoglycan and LTA synthesis.

## Introduction

*Staphylococcus aureus* is a Gram-positive bacterium found as commensal on the skin and nasal passages of healthy individuals. However, this bacterium is also an important human pathogen causing hospital- as well as community-acquired infections, such as serious skin infections, osteomyelitis and endocarditis (1-3). Of major concern is the increasing resistance of this organism to a large number of clinically relevant antibiotics (4). A number of virulence factors contribute to the successful host colonization, immune evasion and acquisition of nutrients within the host (5). Many of these factors are either secreted proteins or other extracellular proteins closely associated with the bacterial cell envelope (6).

The cell envelope is essential for bacterial survival and pathogenesis but also a target of a number of important antimicrobials. It functions as barrier and protects bacteria from environmental insults but at the same time needs to allow the passage of solutes and nutrients as well as sense changes in the external environment (7). *S. aureus* has a typical Gram-positive cell envelop, which consists of a cytoplasmic membrane that is surrounded by a thick peptidoglycan layer (8). The peptidoglycan layer is a dynamic macromolecular structure that undergoes constant cycles of polymerisation and hydrolysis to allow bacteria to grow and divide (7). It is composed of glycan chains made of alternating N-acetylglucosamine and N-acetylmuramic acid residues connected by peptide bridges (9). This mesh-like sacculus is able to protect the cell from environmental threats and at the same time withstand the high internal osmotic pressure (10). The final steps of peptidoglycan synthesis are catalyzed by enzymes named penicillin binding proteins (PBPs) and a coordinated action of these enzymes is crucial for the survival of the cell (11). PBPs with glycosyltransferase and transpeptidase activity polymerize the glycan chains and form peptide cross-bridges while the monofunctioal transpeptidases only have the former activity (11).

*aureus* encodes four PBPs, of which PBP1, which has transpeptidase activity, and the bifunctional PBP2, which has transpeptidase and glycosltransferase activity, are the minimal requirement for cell survival (12). In methicillin-resistant *S. aureus* (MRSA) strains, the alternative PBP2A, which has transpeptidase activity, is needed for β-lactam resistance in addition to the glycosyltransferase activity of PBP2 (13, 14). Moreover, additional non-essential proteins have been identified that are involved in peptidoglycan synthesis such as the monofunctional glycosyltransferases SgtA and SgtB (also named Mgt) (12). Previous studies have shown that although SgtA and SgtB have glycosyltransferase activity *in vitro*, only SgtB can support the growth of *S. aureus* in the absence of the main glycosyltransferase PBP2. However, this is not possible in the presence of β-lactam antibiotics when an interaction between PBP2 and PBP2A is needed (15, 16).

In Gram-positive bacteria, the peptidoglycan layer is interspersed by a plethora of proteins and cell wall polymers named teichoic acids (17). Teichoic acids are further categorised into lipoteichoic acids (LTA), anchored to the outer leaflet of the cytoplasmic membrane via a lipid moiety, and wall teichoic acids (WTA) covalently attached to the peptidoglycan (18). Teichoic acids form an important part of the cell wall and contribute to the physical and chemical properties of the cell wall, and to the binding of divalent cations (19, 20). While both WTA and LTA are polyanioinic cell wall polymers, they are synthesized through separate and independent pathways in *S. aureus* and many other Gram-positive bacteria (21, 22). Consistent with this, our work on the Gram-positive pathogen *Listeria monocytogenes* has revealed that LTA synthesis is not abrogated in the absence of WTA and *vice versa* (23). Recent work using pathway specific inhibitors and a gene interaction screen provided further evidence that the polymers are not only synthesized through separate pathways but also have distinct functions in *S. aureus* (24).

LTA is an anionic polymer made up in *S. aureus* of glycerolphosphate repeating units that are further decorated with D-Alanine residues and, as shown recently, under high salt conditions also with N-acetylglucosamine residues (25, 26). Most proteins required for its synthesis have now been identified and extensively studied over the years (22, 27). One of the key enzymes required for LTA synthesis is the lipoteichoic acid synthase enzyme LtaS (28, 29). This enzyme polymerizes the LTA backbone chain on the outside of the cell using the glycerolphosphate head group of the membrane lipid phosphatidylglycerol as substrate (29, 30). LTA is indispensable for the growth of *S. aureus* under standard laboratory growth conditions, highlighting its important physiological role (29, 31).

Previous studies indicated a function for LTA in helping to direct the cell-division machinery (32), in controlling autolysin activity (31), for biofilm formation (33), in mediating interactions with host cell receptors (17), in controlling susceptibility and/or resistance to antimicrobial peptides and in maintaining cation homeostasis (20). *S. aureus ltaS* mutants, which lack the complete LTA polymer, can in some strain backgrounds be constructed at low growth temperature (31). It has also been shown that LTA-deficient *S. aureus* strains are viable when grown under osmotically stabilizing conditions in medium containing 7.5% NaCl or 40% sucrose (31, 34). However, the bacteria display severe morphological defects including an enlarged cell size, clustering and cell division defects, even under those conditions permissive for growth (34). Bacteria can also readily acquire compensatory mutations allowing them to grow in un-supplemented medium and improving their cell morphology defects (34). The majority of compensatory mutations previously observed were in *gdpP* leading to inactivation of the c-di-AMP phosphodiesterase GdpP (34). The resulting increase in cellular c-di-AMP levels allowed the bacteria to survive the cell wall stress caused by the absence of LTA, which is now believed to be due to changes in the osmotic balance in the cell (34). However, compensatory mutations that rescue the growth of an LTA negative *S. aureus* strain were not only found in *gdpP* but also in other genes (34).

As part of the current study, we sought to identify these genes and further characterize the encoded proteins to gain additional insight into why LTA is essential for the growth of *S. aureus* and potentially uncover novel cellular functions for proteins involved in cell wall assembly or maintenance. Using a suppressor screen approach, we found that inactivation of the monofunctional peptidoglycan glycosyltransferase SgtB allows *S. aureus* to grow in the absence of LTA. We further show that peptidoglycan crosslinking is significantly reduced in the absence of LTA, which can be partially restored upon inactivation of SgtB. This in turn might strengthen the peptidoglycan layer and, in this way, contribute to the observed growth rescue.

## Results

### Identification of *S. aureus* suppressor strains able to grow in the absence of LTA in a c-di-AMP-independent way

LTA-deficient *S. aureus* strains are viable when grown under osmotically stabilizing conditions in medium containing 7.5% NaCl or 40% sucrose or in un-supplemented medium after the acquisition of compensatory mutations (31, 34). The majority of compensatory mutations previously observed were in *gdpP* leading to inactivation of the c-di-AMP phosphodiesterase GdpP (34). In the same study, suppressor strains with mutations outside the *gdpP* gene were noted (34). In order to characterize suppressor strains with mutations in genes other than *gdpP* and to gain further insight into the cellular function of LTA, a larger suppressor screen was performed and the *ltaS* mutant strains LAC*Δ*ltaS∷erm* constructed in medium supplemented either with sucrose or NaCl were plated on un-supplemented TSA plates. A number of independently obtained suppressor colonies were subsequently passed 4 times in fresh TSB to further improve their growth. Next, the chromosomal DNA was isolated from 80 suppressor strains and those lacking mutations in *gdpP* (coding for the c-di-AMP hydrolase) and *dacA* (coding for the c-di-AMP cyclase enzyme) were identified by determining the sequences of these two genes. Of 80 colonies screened, none had mutations in *dacA* and 17 strains had no mutation in *gdpP*, 7 of which were selected for further analysis. The absence of LTA but presence of WTA in the suppressor strains was confirmed by western blot and Alcian blue silver staining analysis, respectively (Figs. 1A and 1B). Next, the relative cellular c-di-AMP levels in the different strains were determined using a previously described competitive ELISA assay (35, 36) (Fig. 1C). In contrast to the *gdpP* mutant or LTA negative suppressor strain with a mutation in *gdpP*, which showed the expected increase in c-di-AMP levels, the 7 suppressor strains chosen for further analysis did not show an increase in the cellular c-di-AMP concentration (Fig. 1C).

**Figure 1:**
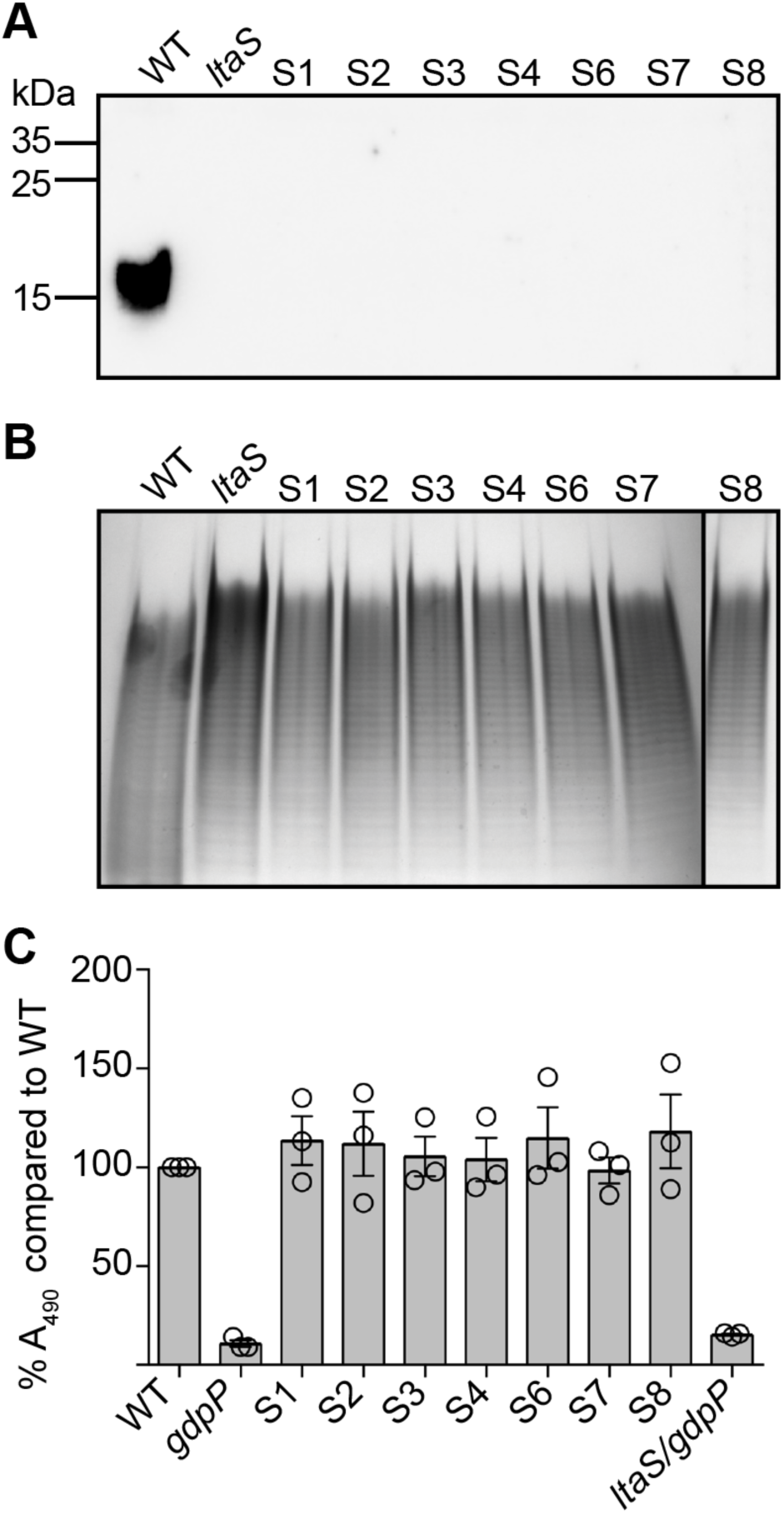
Detection of LTA by western blot, WTA by Alcian blue silver staining and c-di-AMP using a competitive ELISA method. (A) Detection of LTA by western-blot. Cell extracts were prepared from LAC* (WT), an *ltaS* mutant (strain ANG2135) and the 7 *ltaS* suppressor strains (S1-S4 and S6-S8) and separated on a 15% PAA gel. LTA was subsequently detected by western blot using a monoclonal polyglycerolphosphate specific antibody. A representative result from 3 independent experiments is shown. (B) Visualization of wall teichoic acid (WTA) by native PAGE. WTAs were isolated from the same strains as shown in (A) and separated on a 20% native gel. Bands were visualized by Alcian blue and subsequently silver stain. One representative result from 4 independent experiments is shown. (C) c-di-AMP detection by competitive ELISA. Cytoplasmic extracts were prepared from LAC* (WT), the high c-di-AMP level control strains LAC* *gdpP∷kan* (*gdpP*) and US3 (*ltaS/gdpP*) as well as from the 7 *ltaS* suppressor strains (S1-S4 and S6-S8). c-di-AMP amounts were determined and compared to the WT strain by ELISA. Of note, as this is a competitive ELISA assay, lower A_490_ readings are obtained for samples with higher c-di-AMP levels. The A_490_ reading obtained for the sample derived from the WT strain was set to 100% and % A_490_ reading calculated for the test strains. The average % values and standard deviations from three independent experiments (with three technical replicates) are plotted.

### Identification of genomic alterations in the *S. aureus* suppressor strains able to grow in the absence of LTA

Next, the genomic sequences of the seven suppressor strains were determined and compared to that of the original *ltaS* mutant strains using a whole genome sequencing approach. Mutations were found in *sgtB* (SAUSA300_1855), coding for the monofunctional glycosyltransferase SgtB and mutations in this gene arose independently in three suppressor strains (Table 1). Another strain had a mutation in SAUSA300_1254, coding for a hypothetical membrane protein as well as SAUSA300_RS11150, encoding MazE, the antitoxin component of a type II toxin-antitoxin module (Table 1). In the original study by Corrigan *et al*. (34), mutations found in SAUSA300_1254 were proposed to be accessory and required to further improve the growth of the *gdpP* mutant suppressor strains. Consistent with a previous report (2), a large deletion in *clpX* (SAUSA300_1621), encoding a protein forming part of an ATP-dependent protease, was observed in one strain (Table 1). Finally, a mutation in *vraT* (SAUSA300_1867), coding for the membrane protein VraT and forming part of the VraRST three-component system, was identified. Using our standard genome sequence analysis workflow, an unusually large number (>300) of zero coverage regions were obtained for suppressor strain S6, preventing us from matching a single nucleotide polymorphism with high confidence to the observed growth rescue. Hence, suppressor stain S6 is not listed in Table 1 and was also not further analyzed. The mutations identified in the other suppressor strains were subsequently confirmed by fluorescent automated resequencing of the respective genomic region. Some of the mutations observed in *sgtB*, as well as the mutations in *vraT, clpX* and *mazE* result in frameshift mutations and introduction of premature stop codons, suggesting that the absence of the encoded proteins compensates for the lack of LTA.

**Table 1:**
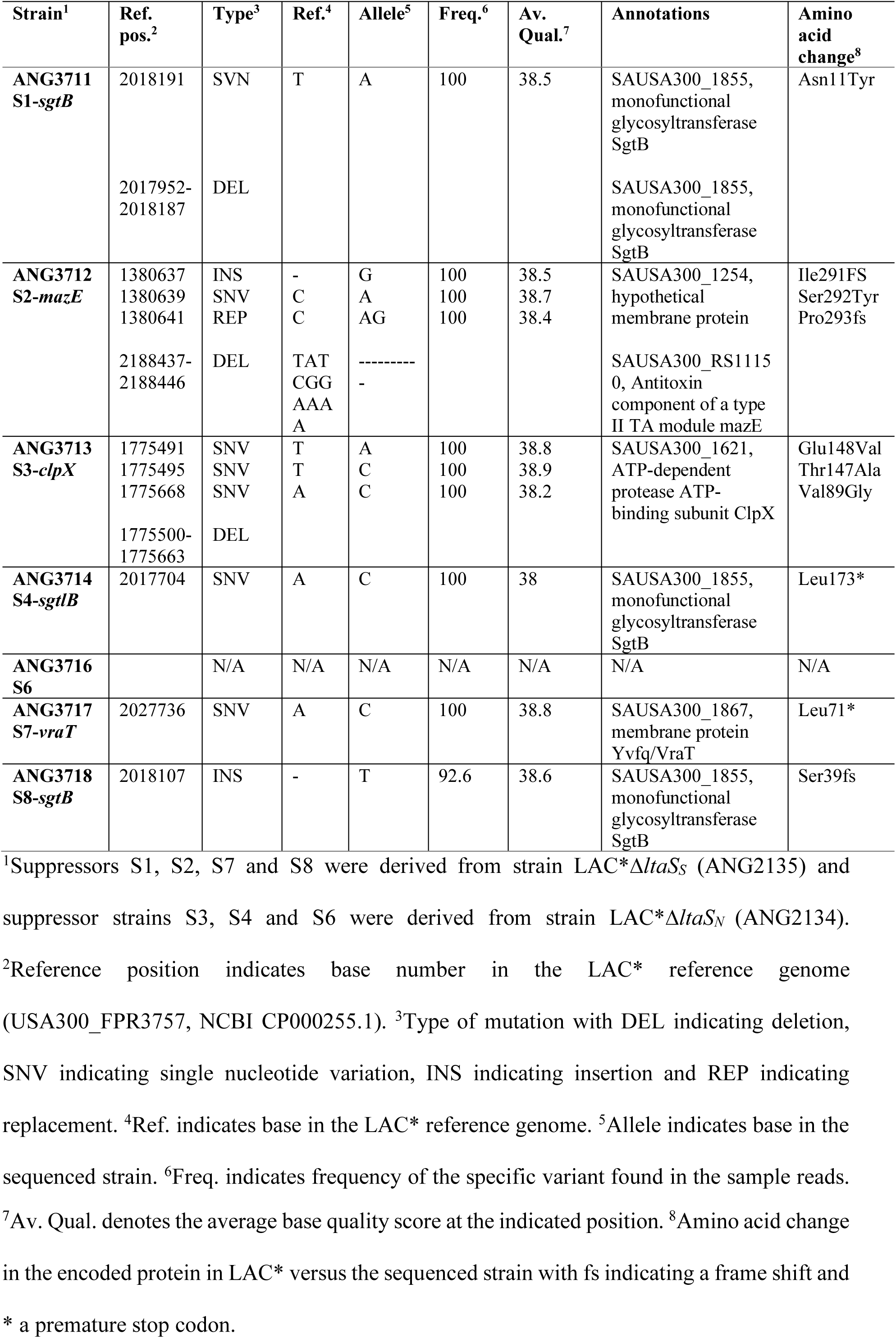
Genomic variations detected in the suppressor strains as compared the original *ltaS* mutant strains

### Phenotypic characterization of the LTA negative *S. aureus* suppressor strains

In a previous study, it was shown that *S. aureus* cells can grow without LTA in the absence of ClpX (2), hence we did not further characterize the *clpX* mutant suppressor strain obtained as part of this study. Instead our further analysis focused on suppressor strains with mutations in novel genes, which are strains with mutations in *mazE, sgtB* or *vraT*. MazE, the antitoxin component of a type II TA module, is part of the *sigB* regulon and has been shown to be essential for full activity of the alternative sigma factor SigB (37). SigB and members of its regulon enable bacteria to respond rapidly to environmental and antibiotic stresses and also play a role in cell envelope homeostasis (38, 39). Various studies have investigated the role of the VraTSR three-component regulatory system and this system has been reported to be involved in the induction of the cell wall stressosome, mainly in the presence of cell wall targeting antibiotics (40-42).

Interestingly, the monofunctional glycosyltransferase SgtB, also identified as part of our screen, is one of the proteins belonging to the cell wall stressosome whose expression is regulated by VraTSR independent of the presence of cell wall targeting antibiotics (40). To further characterize the suppressor strains, we first confirmed the growth improvement of strains S2-*mazE*, S4-*sgtB* and S7-*vraT* when propagated in TSB. All three suppressor strains grew better compared to the original *ltaS* mutant strain and their growth rate was only slightly reduced compared to the WT LAC* stain (Fig. 2A). Next, the cell morphology of the WT, original *ltaS* mutant and the three suppressor strains was assessed by microscopy following staining of the peptidoglycan with fluorescently labelled vancomycin. The cell morphology of the suppressor strains was considerably improved and in particular for suppressor strain S2-*mazE* the division site was correctly placed in most cells (Fig. 2B). Next, the susceptibility of the suppressor strains to a number of cell wall targeting antibiotics was determined. Although the growth and cell morphology of the suppressor strains were improved, these strains remained hyper-sensitive to the β-lactam antibiotic oxacillin and the MIC was reduced by 32-fold or more for the different suppressor strains as compared to the WT LAC* strain (Fig. 3A). The susceptibility to the cell wall or membrane targeting antibiotics lysostaphin, nisin, vancomycin or daptomycin was also tested but no drastic differences were observed (Table 2). Finally, the growth of the three different suppressor strains was assessed on plates containing Congo red. Congo red is an anionic azo dye traditionally used for the detection of biofilms in *Staphylococcus*. However, at higher concentrations it inhibits the growth of *S. aureus* and has been used in the past to indicate differences in the cell wall integrity of different *S. aureus* strains (43). Recently, the target of Congo red has been established as the LTA synthase enzyme LtaS (44). Hence our suppressor strains, which are deleted for *ltaS* and are able to grow in the absence of LTA, should be resistant to this dye. To test the susceptibility of the suppressor strains, serial dilutions of overnight cultures were spotted on TSA plates containing 0.1% Congo red (Fig. 2C). The suppressor strains carrying mutations in *mazE* and *sgtB* were significantly more resistant to Congo red than the WT strain, indicating that inactivation of either one of these genes is indeed sufficient to bypass the LTA essentiality. However, the suppressor strain with the mutation in *vraT* grew poorly on the Congo red plates, suggesting that this strain might not be a bona fide suppressor strain.

**Table 2:** MIC values for different antibiotics given in µg/ml

|  | **lysostaphin** | **nisin** | **vancomycin** | **daptomycin** |
| --- | --- | --- | --- | --- |
| **LAC\*** | **0.125**<br>(0.125, 0.125, 0.125) | **16**<br>(16, 16, 16) | **4**<br>(4, 4, 4) | **16**<br>(16, 16, 16) |
| **S2-mazE** | **0.125**<br>(0.125, 0.125, 0.125) | **2-4**<br>(2, 2, 4) | **2**<br>(2, 2, 2) | **4-8**<br>(8, 8, 4) |
| **S4-*sgtB*** | **0.125-0.250**<br>(0.125, 0.125, 0.25) | **4**<br>(4, 4, 4) | **1**<br>(1, 1, 1) | **8-16**<br>(16, 8, 8) |
| **S7-*vraT*** | **0.500**<br>(0.5, 0.5, 0.5) | **2-4**<br>(2, 2, 4) | **4-8**<br>(8, 4, 4) | **8**<br>(8, 8, 8) |
| ***sgtB::tn*** | **0.125-0.250**<br>(0.25, 0.125, 0.125) | **16-32**<br>(32, 16, 32) | **4**<br>(4, 4, 4) | **16**<br>(16, 16, 16) |
| ***sgtB::tn***<br>**pCL55** | **0.125-0.250**<br>(0.125, 0.125, 0.25) | **8-16**<br>(16, 8, 16) | **4**<br>(4, 4, 4) | **16**<br>(16, 16, 16) |
| ***sgtB::tn***<br>**pCL55-*sgtB*** | **0.125**<br>(0.125, 0.125, 0.125) | **8-16**<br>(8, 16, 16) | **4-8**<br>(4, 8, 4) | **16**<br>(16, 16, 16) |

**Figure 2:**
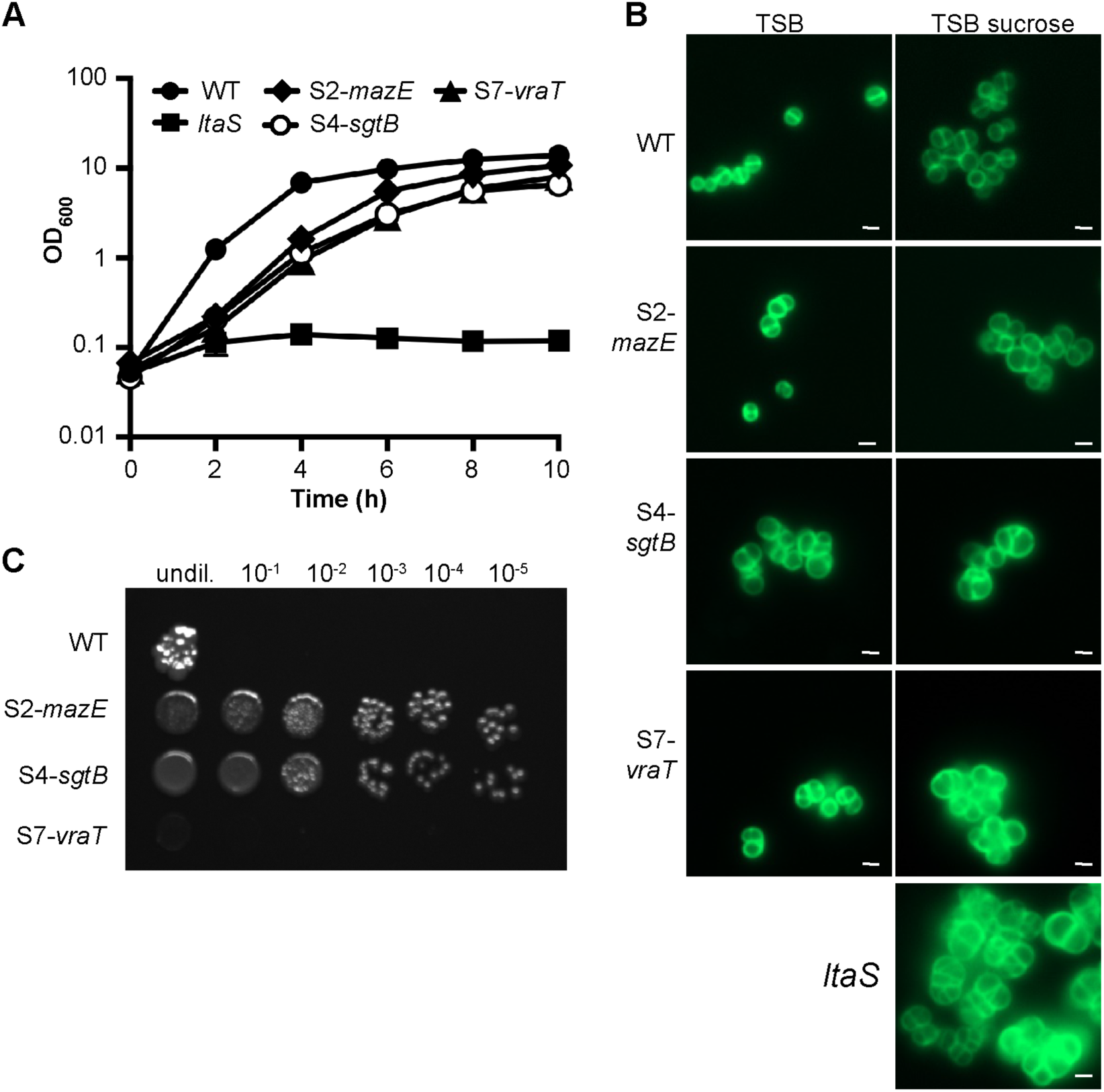
Growth and cell morphology of WT and mutant *S. aureus* strains. (A) Bacterial growth curves. LAC* (WT) and the suppressor strains S2*-mazE,* S4-*sgtB* and S7-*vraT* were grown overnight in TSB and the original *ltaS* mutant strain (ANG2135) in TSB with 40% sucrose. Next day, bacterial cells were washed and diluted in TSB to an OD_600_ of 0.05 and the bacterial growth was subsequently monitored over a period of 10 hours. The average OD_600_ readings from three experiments were plotted. (B) Microscopic analysis. Bacterial cells from overnight cultures of the same strains as used in panel A were washed, back-diluted and grown to mid-log phase in TSB or TSB with 40% sucrose as indicated. The bacterial cells were subsequently stained with BODIPY-vancomycin and viewed by fluorescent microscopy. (C) Analysi*s* of bacterial growth on Congo red-containing TSA plates. Overnight cultures of WT and the indicated suppressor strains were serially diluted and spotted on TSA plates supplemented with 0.1% Congo red and the plates incubated for 48h at 37°C. For B and C, a representative result of three independent experiments is shown.

**Figure 3:**
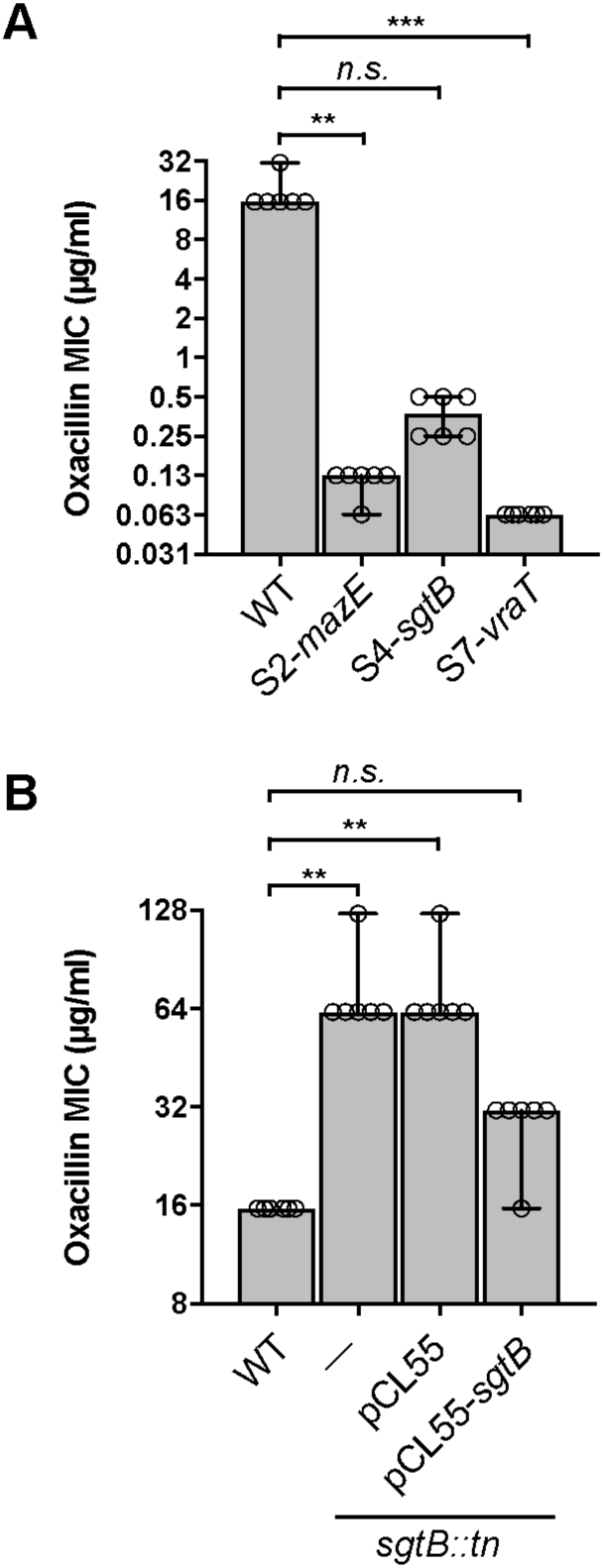
Oxacillin MICs for WT and mutant *S. aureus* strains. Oxacillin minimal inhibitory concentrations (MICs) of (A) LAC* (WT) and the suppressor strains S2*-mazE,* S4-*sgtB* and S7-*vraT* and (B) LAC* (WT), the LAC* *sgtB∷tn* mutant either without integrative plasmid (-), with integrative plasmid (pCL55) or integrative complementation plasmid (pCL55-*sgtB*). Experiments were conducted with six biological replicates. Medians and 95% confidence intervals for all strains are plotted. A Kruskal-Wallis one-way analysis of variance tests was performed, which indicated significant differences. Subsequently, Dunn’s tests were performed and the p-values corrected for the multiple comparisons against the WT. p-values < 0.01 (**) and < 0.001 (***).

### Introduction of SgtB or MazE in the respective suppressor strain results in the expected growth arrest

The result of the whole genome sequencing analysis suggested that inactivation of MazE or SgtB is sufficient to allow *S. aureus* to grow in the absence of LTA. Introduction of a wild-type copy of *mazE* or *sgtB* into the respective suppressor strain should restore this phenotype and be lethal for the suppressor strains when grown in un-supplemented medium but should not have an effect when the bacteria are propagated in medium supplemented with 40% sucrose. In order to test this, plasmids piTET-*mazE* or piTET-*sgtB,* allowing for anhydrotetracycline (Atet) inducible gene expression, were introduced into the respective suppressor strains. As control, these plasmids were also introduced in the WT LAC* strain. Serial dilutions of these different strains were spotted onto TSA plates containing 100 ng/ml or 200 ng/ml Atet for the expression of *mazE* or *sgtB* expression, respectively. As expected for a successful complementation, the expression of *mazE* prevented the growth of the suppressor strain S2-*mazE* (Fig. 4A) and the expression of *sgtB* prevented the growth of the suppressor strain S4-*sgtB* (Fig. 4B) on TSA but had no effect when the bacteria were spotted on medium supplemented with 40% sucrose (Fig. 5). Of note, the *mazE* suppressor strain S2-*mazE* contained an additional mutation in a gene coding for a membrane protein of unknown function with locus tag SAUSA300_125. Introduction of a WT copy of this gene in the suppressor strain did not prevent the growth of this suppressor strain (data not shown). Taken together, the results of this complementation analysis support the notion that inactivation of SgtB or MazE is sufficient to allow *S. aureus* to grow in the absence of LTA.

**Figure 4:**
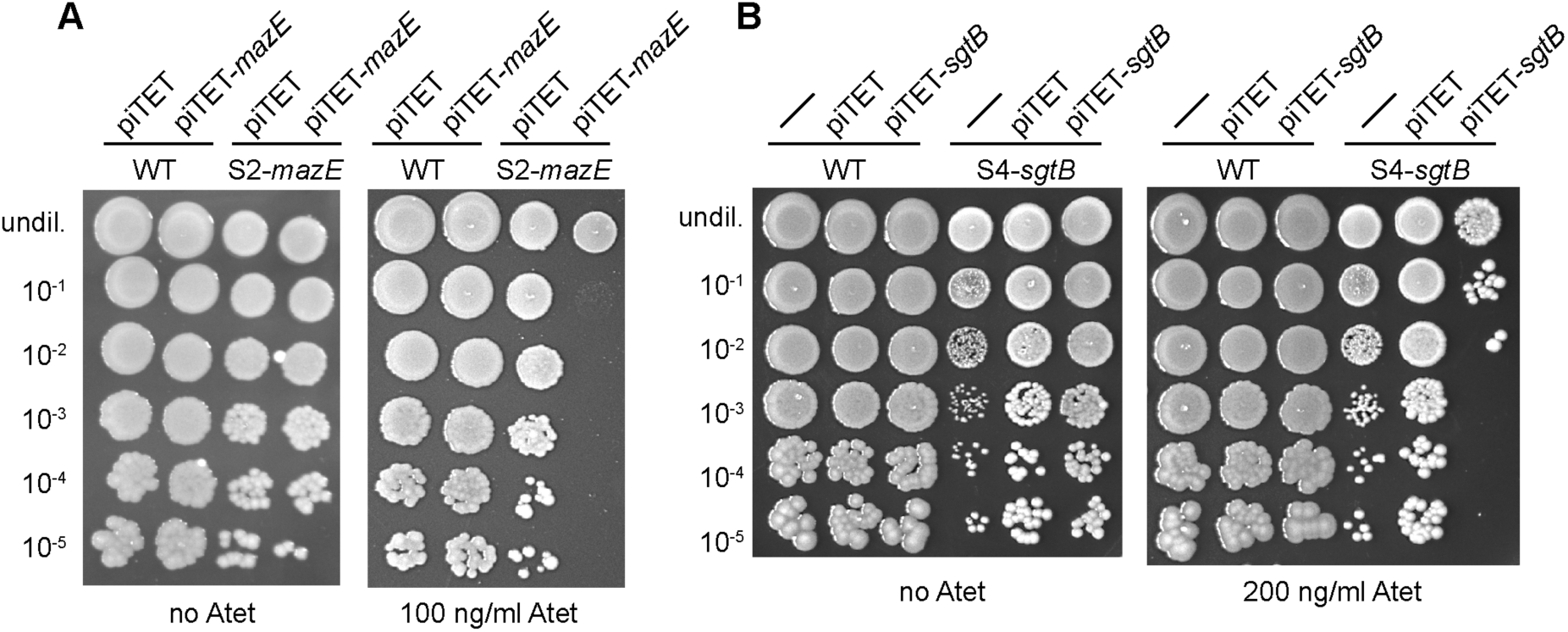
Growth complementation analysis. TSA plates with spot dilutions. (A) *S. aureus* strains LAC* piTET, LAC* piTET-*mazE*, S2-*mazE* piTET or S2-*mazE* piTET-*mazE* were grown overnight in TSB supplemented with chloramphenicol, washed twice in PBS and serial dilutions spotted on TSA plates supplemented with 7.5 ng/ml chloramphenicol without (left panel) or with 100 ng/ml Atet (right panel). (B) *S. aureus* strains LAC* (WT), LAC* piTET, LAC* piTET-sgtB, S4-*sgtB*, S4-*sgtB* piTET or S4-*sgtB* piTET-*sgtB* were grown and samples prepared as described in A, however without chloramphenicol selection, and dilutions were spotted on TSA plates (left panel) or TSA plates supplemented with 200 ng/ml Atet (right panel). Representative plate images from three independent experiments are shown.

**Figure 5:**
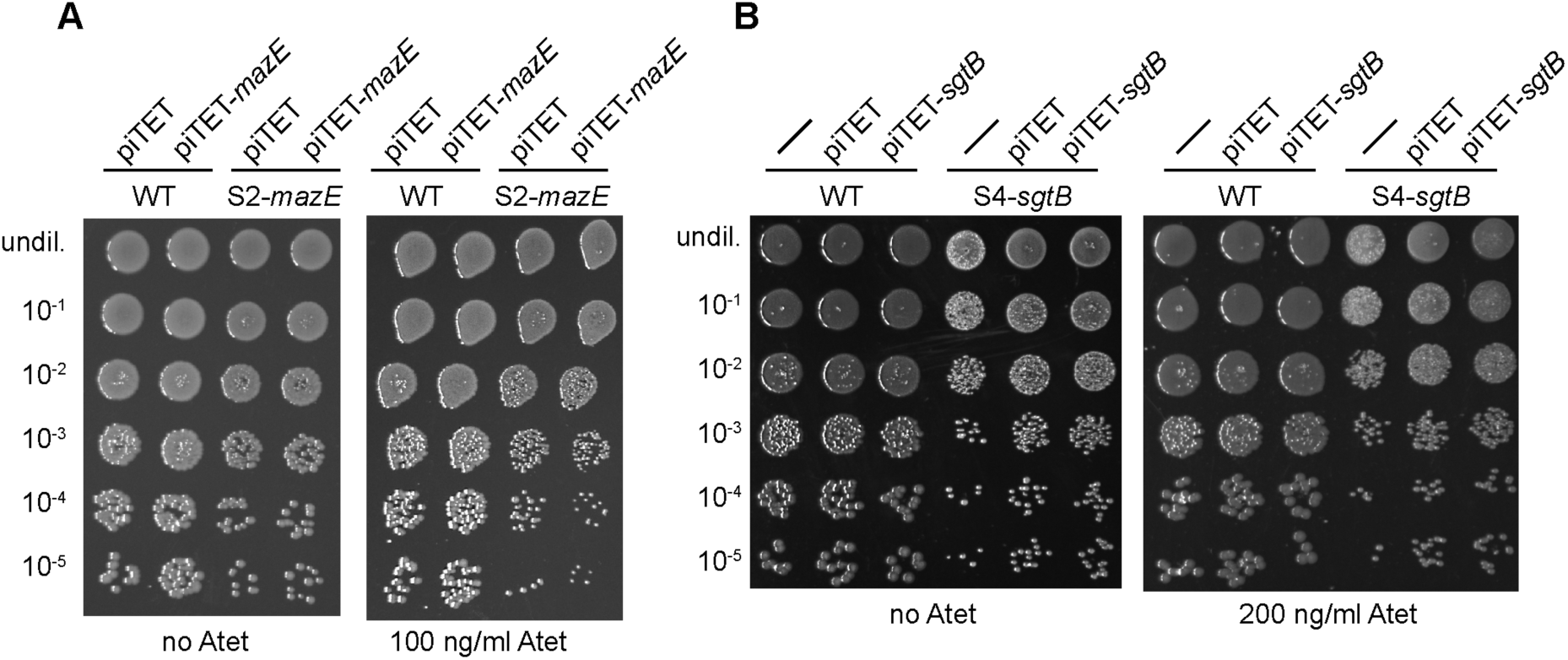
Growth complementation analysis. TSA sucrose plates with spot dilutions. (A) *S. aureus* strains LAC* piTET, LAC* piTET-*mazE,* S2-*mazE* piTET or S2-*mazE* piTET-*mazE* were grown overnight in TSB supplemented with chloramphenicol, washed twice in PBS and serial dilutions spotted on TSA 40% sucrose plates supplemented with 7.5 ng/ml chloramphenicol without (left panel) or with 100 ng/ml Atet (right panel). (B) *S. aureus* strains LAC* (WT), LAC* piTET, LAC* piTET-*sgtB,* S4-*sgtB,* S4-*sgtB* piTET or S4-*sgtB* piTET-*sgtB* were grown and samples prepared as described in A, however without chloramphenicol selection, and dilutions were spotted on TSA 40% sucrose plates (left panel) or TSA 40% sucrose plates supplemented with 200 ng/ml Atet (right panel). Representative plate images from three independent experiments are shown.

### Growth characterization and antibiotic resistance of an *S. aureus* LAC* *sgtB* mutant

Inactivation of MazE likely has pleotropic effects due to its involvement in the activity of the alternative sigma factor SigB and the reason as to why its inactivation allows *S. aureus* to grow in the absence of LTA could be indirect. Hence, we next focused in trying to gain a better understanding of the cellular function of the monofunctional peptidoglycan glycosyltransferase SgtB and how its inactivation allows *S. aureus* to survive in the absence of LTA. To this end, the *sgtB* mutant strain LAC\**sgtB∷tn* was constructed by moving the genomic region with a transposon insertion in *sgtB* from the Nebraska transposon mutant library strain NE596 (45) into the LAC* background. SgtB is one of two monofunctional peptidoglycan glycosyltransferases encoded in the *S. aureus* genome and the protein can polymerize peptidoglycan glycan chains *in vitro* (46). Although it is dispensable for the growth of *S. aureus*, SgtB becomes necessary for bacterial survival in the absence of the main glycosyltransferase PBP2 (12, 15, 47). Consistent with these previous observations, no difference was observed in the cell growth and morphology of the *sgtB* mutant *S. aureus* strain LAC\**sgtB∷tn* as compared to the WT LAC* strain (Fig. 6A and 6B). Our results indicate that inactivation of SgtB allows *S. aureus* to survive in the absence of LTA, therefore the *sgtB* mutant strain should no longer be sensitive to the Congo red dye. To test this, serial dilutions of the WT, the *sgtB* mutant and a complementation strain were spotted on TSA plates containing 0.1% Congo red, which inhibits the LtaS enzyme. Indeed, the *sgtB* mutant strain was considerably more resistant to Congo red than the WT strain and this phenotype could be complemented by introducing a functional copy of *sgtB* into the mutant strain (Fig. 6C). Next, we also tested the susceptibility of the *sgtB* <u>mutant and complementation strain towards the cell wall active antibiotics</u> oxacillin, lysostaphin, nisin, vancomycin and daptomycin. While no differences were observed in the susceptibility of the *sgtB* mutant compared to a wild-type strain towards most antibiotics, a slight (4-fold) and statistically significant increase in resistance towards oxacillin was observed (Fig. 3B, Table 2). Taken together, while no growth, or clearly visible morphological differences were observed between the WT and the *sgtB* mutant strain, deletion of *sgtB* leads to increased Congo red dye and 4-fold increased oxacillin antibiotic resistance in our strain background.

**Figure 6:**
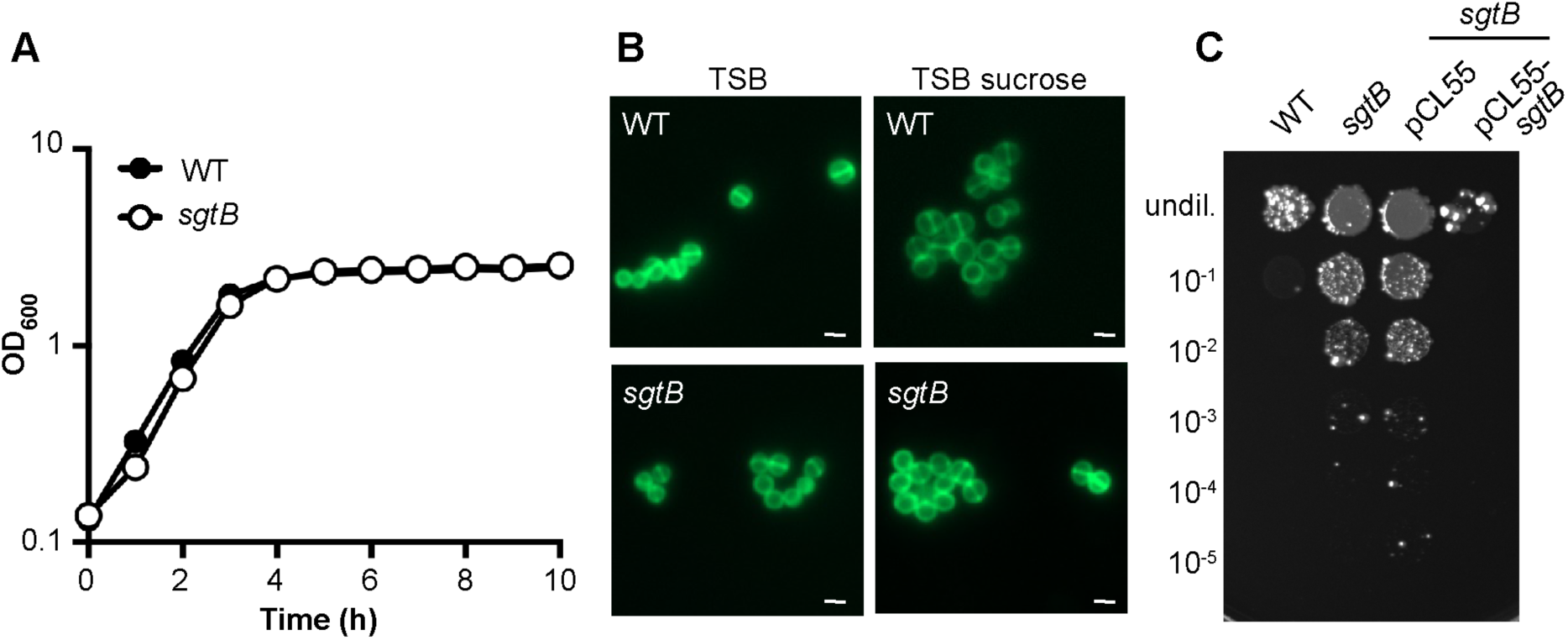
Growth and cell morphology of *S. aureus* strain LAC\**sgtB∷tn*. (A) Bacterial growth curves. Overnight cultures of LAC* (WT) and strain LAC\**sgtB∷tn* (*sgtB*) were diluted in TSB to an OD_600_ of 0.05 and the growth monitored over a period of 10 hours using a plate reader. The average OD_600_ values and standard deviation from three independent experiments are plotted. (B) Microscopy analysis. *S. aureus* strains LAC* (WT) and LAC\**sgtB∷tn* (*sgtB*) were grown overnight in TSB or TSB supplemented with 40% sucrose, stained with BODIPY-vancomycin and viewed under a fluorescence microscope. (C) Susceptibility to Congo red. Overnight cultures of strains LAC* (WT) and the *sgtB* mutant strains LAC\**sgtB∷tn*, LAC\**sgtB∷tn* pCL55 as well as the complementation strain LAC\**sgtB∷tn* pCL55-*sgtB* were serially diluted and aliquots spotted on TSA plates supplemented with 0.1% Congo red. For panels B and C, representative result from three independent experiments are shown.

### Inactivation of SgtB leads to an increase in peptidoglycan cross-linking in an LTA-negative *S. aureus* strain

Since SgtB is involved in peptidoglycan synthesis and the *sgtB* mutant displayed increased slightly increased oxacillin resistance, we hypothesised that its deletion could somehow “strengthen” the cell wall through alterations in the peptidoglycan structure. In order to investigate this, the muropeptide profiles of peptidoglycan isolated from the WT LAC*, the *sgtB* mutant strain LAC\**sgtB∷tn*, and the *ltaS/sgtB* mutant suppressor strain S4-*sgtB* were determined following the growth of these strains in TSB. In addition, the muropeptide profiles were also determined for peptidoglycan isolated from these three strains as well as the original *ltaS* mutant strain following growth in TSB supplemented with 40% sucrose (Fig. 7). The WT and *sgtB* mutant strains showed very similar and typical *S. aureus* muropeptide profiles (Fig. 7A). The chemical structure of several muropeptide fragments for a number of these peaks has been previously determined in the seminal paper by de Jonge *et al.* (48). We numbered the different peaks where possible as described in de Jonge *et al.* (48) with peaks 1-5 being monomeric, 9–14 dimeric, 15 trimeric and 16 and above higher oligomeric muropeptide fragments. Quantification of the monomeric, dimeric, trimeric and higher oligomeric peptidoglycan fragments showed that the peptidoglycan is highly crosslinked in both strains with approximately 70% of the UV absorbing material found in the higher oligomeric fraction (Fig. 7B and 7C). No clear differences were found between the muropeptide profiles of the WT and *sgtB* mutant strain, either grown in TSB or TSB supplemented with 40% sucrose (Fig. 7). In contrast to the WT and *sgtB* mutant strain, a decrease in the higher-oligomeric peptidoglycan material was observed for the *ltaS* suppressor strain S4-*sgtB* both in TSB and TSB sucrose medium (Fig 7). A visual inspection of the chromatograms also indicated that peak 12, corresponding to a currently unknown muropeptide, was reduced in *S. aureus* strains unable to produce LTA, which are the original *ltaS* mutant as well as the *S4-ltaS* suppressor strain (Fig. 7A). But perhaps most notably, a comparison of the muropeptide profiles of the peptidoglycan isolated from the original *ltaS* mutant and the *ltaS/sgtB* suppressor strain S4-*sgtB* after growth in TSB sucrose medium showed that, while the amount of crosslinked peptidoglycan was reduced in both strains compared to the WT, the peptidoglycan in the suppressor strain was more crosslinked as compared to the original *ltaS* mutant strain (Fig 7A and 7B). Taken together, our data highlight that deletion of *ltaS* leads to a significant reduction in the amount of crosslinked peptidoglycan in *S. aureus*. Inactivation of SgtB in a WT strain does not significantly affect the amount of crosslinked peptidoglycan, as one might have expected based on the observed increase in resistance oxacillin. However, in an LTA negative background strain, inactivation of SgtB leads to an increase in peptidoglycan crosslinking, which might explain why an LTA negative strain can grow in the absence of this monofunctional peptidoglycan glycosyltransferase.

**Figure 7:**
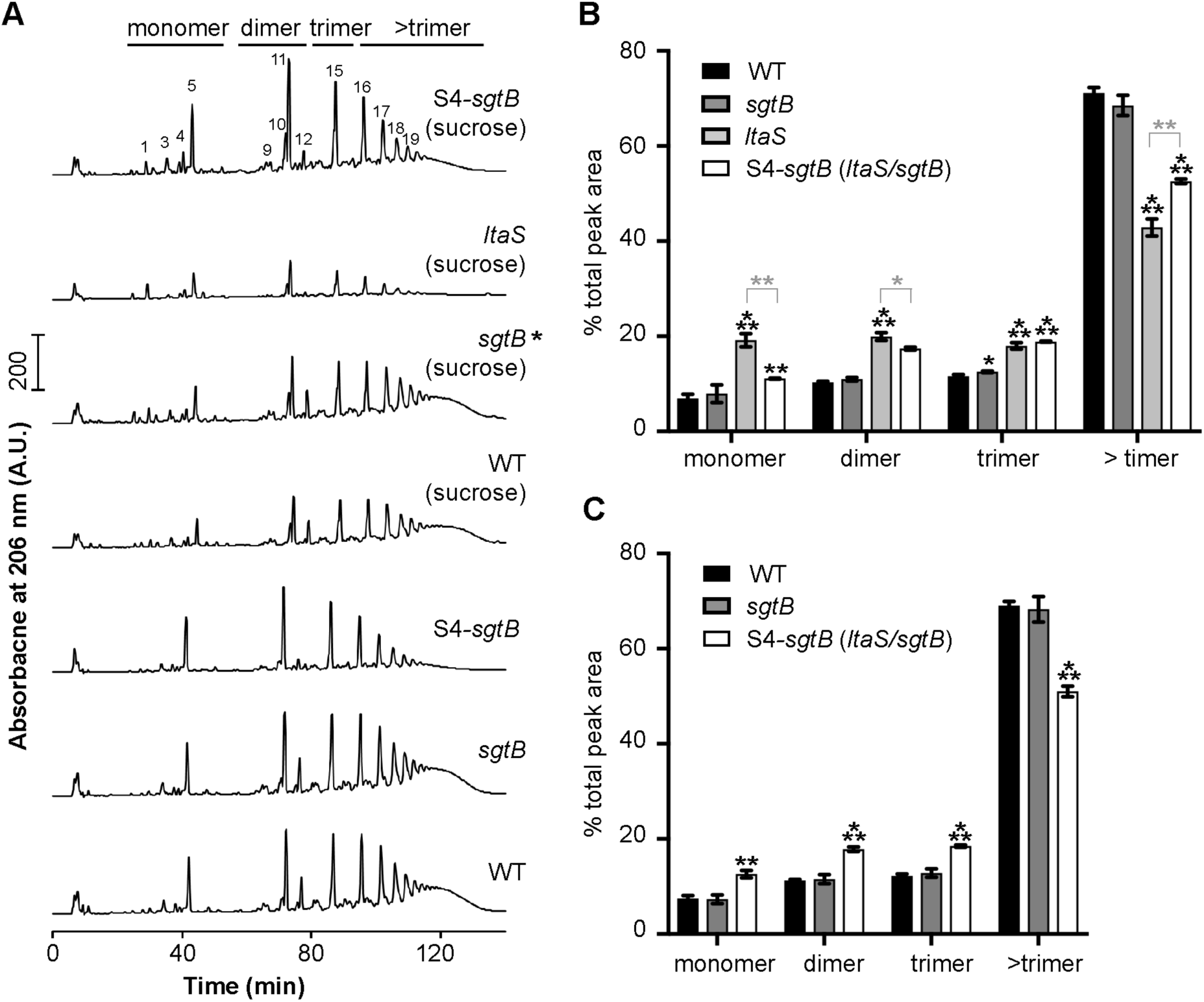
Peptidoglycan analysis of WT and mutant *S. aureus* strains. (A) HPLC profile of mutanolysin digested peptidoglycan. Peptidoglycan was isolated and digested with mutanolysin as described in the materials and methods section from *S. aureus* strains LAC* (WT), LAC\**sgtB∷tn* (*sgtB*) and the LTA negative suppressor strain S4-*sgtB* following growth in TSB medium or from *S. aureus* strains LAC* (WT), LAC\**sgtB∷tn* (*sgtB*), the LTA negative suppressor strain S4-*sgtB* as well as the original *ltaS* mutant strain (ANG2135) following growth in TSB 40% sucrose medium. Monomeric, dimeric, trimeric and > trimeric peptidoglycan fragments are indicated above the graphs and a representative profile from three independent samples is shown. The individual muropeptide peaks were labelled where possible with the numbers as described in de Jonge *et al*. (48). (B-C) Quantification of the different peptidoglycan peaks. The peaks corresponding to monomeric, dimeric, trimeric and > trimeric peptidoglycan fragments were integrated and quantified for the strains used in panel A. The combined peak area was set to 100% for each strain and the average values and standard deviations from the three independent peptidoglycan isolations were plotted in panel B, for the strains grown in TSB sucrose medium and in panel C for the strains grown in TSB medium. Unpaired two-tailed student’s T-tests were used to determine statistically significant differences in monomer, dimer, trimer or > trimer fractions between WT and mutant strains. The obtained p-values were multiplied by the Bonferroni correction factors 3 (panel B) and 2 (panel C) to consider that the WT peaks were compared to three or two different strains, respectively. Statistically significant differences are indicated by black asterisks. For panel B, unpaired two-tailed student’s T-tests were also used to determine statistically significant differences in monomer, dimer, trimer or > trimer fractions between the *ltaS* mutant and the other three strains and the statistically significant differences between the *ltaS* mutant and S4-*ltaS* suppressor are shown with grey asterisks. p-values < 0.05 (*), < 0.01 (**) and < 0.001 (***).

## Discussion

The anionic LTA polymer is a core component of the cell wall, essential for survival and a number of studies have shown its importance in various cell processes (see review (22)). Phenotypes caused by the depletion of LTA in *S. aureus* are misplacement and incomplete formation of cell division septa, enlargement of the cells, together ultimately leading to cell lysis (29, 34). However, how LTA mediates these roles is still unknown. In previous work, it has been shown that LtaS and other core LTA synthesis proteins physically interact with early and late stage cell division proteins as well as with a number of peptidoglycan synthesis proteins (49). This indicates that LTA synthesis enzymes might at least transiently be part of multi-enzyme complexes which might help to coordinate LTA synthesis with peptidoglycan synthesis and cell division (49).

We have previously shown that *S. aureus* mutants producing increased intracellular c-di-AMP levels can survive without LTA (34). It is now believed that at high c-di-AMP levels and through the c-di-AMP-dependent regulation of potassium and osmolyte transporters, the internal turgor pressure in the cell might be reduced so that the compromised LTA-depleted cell wall can sustain the internal pressure (34, 50-52). Indeed, as part of the current study we show that the absence of LTA leads to a sizable reduction in the amount of cross-linked peptidoglycan in *S. aureus* (Fig. 7). This is consistent with the idea that in the absence of LTA the cell wall is likely less able to sustain the high internal turgor pressure, which might also contribute to the increased β-lactam sensitivity of LTA-negative strains observed in this and previous studies (Table 2) (2, 34). The reason for the reduced amount of peptidoglycan crosslinking is currently not clear but could potentially be due to mislocalization of PBPs in the absence of LTA or PBPs having reduced enzyme activity since the proper ion homeostasis cannot be maintained within the cell wall in the absence of LTA.

The aim of this study was to further elucidate the role of LTA in cell wall assembly and potentially uncover additional proteins involved in the maintenance of cell wall integrity. A suppressor screen followed by whole genome sequencing revealed mutations in genes coding for ClpX, SgtB, MazE and VraT (Table 1) that can bypass the essentiality of LTA. Further experimentation indicated that the strain with the mutation in *vraT* might not be a bona fide suppressor strain. On the other hand, complementation analysis and growth assays on agar plates containing the azo dye Congo red that prevents the growth of *S. aureus* by inhibiting the LtaS enzyme (44) confirmed that inactivating mutations in *sgtB* and *mazE* could bypass the essentiality of LTA. In a previous study it was found that an *S. aureus clpX* mutant readily acquires inactivating mutations in *ltaS* resulting in the generation of LTA negative strains (2). Conversely and consistent with this previous work, in this study we found in one of our LTA-negative suppressor strains a large deletion in *clpX* (Table 1). ClpX is a protein folding chaperon, which recognizes and targets proteins for degradation to the ClpP protease component. In the absence of ClpX, *S. aureus* cells become smaller, show increased production of autolysins and bacteria have a severe growth defect at temperatures of 30°C or lower (2). The introduction of loss of function mutations in the gene coding for the LTA synthase LtaS in a *clpX* mutant alleviates some of these effects. This is perhaps due to LTA depletion having the opposite effect that leads to an increase in cell size and decreased in autolysis as reported in some previous publications (2, 31).

Mutations in the monofunctional glycosyltransferase SgtB were the most prevalent mutations that arose in our suppressor screen. Previous studies have shown that SgtB is not essential for the survival of the cell and as reported in this and a previous study, an *sgtB* mutant strain does not show any obvious growth or morphological differences as compared to a WT strain under standard growth conditions (Fig. 6) (15). Interestingly, we found that the *sgtB* mutant strain LAC\**sgtB∷tn* was 4-fold more resistant to the cell-wall targeting antibiotic oxacillin and this phenotype could be complemented by introduction of a wild-type copy of *sgtB* (Table 2). These results are in accordance with previous reports, where strains with mutations in genes that compensate for the lack of LTA, such as *gdpP* and *clpX* mutant strains, also show increased oxacillin resistance (2, 34).

The increased resistance to cell wall-targeting antibiotics of the *sgtB* mutant prompted us to investigate the peptidoglycan structure of an *sgtB* mutant in more detail, as we hypothesized that in its absence changes such as an increased crosslinking might be observed that could potentially explain the increased resistance. However, we could not detect any obvious changes in the muropeptide profile of an *sgtB* mutant strain as compared to a WT strain (Fig. 7). On the other hand, the original *ltaS* mutant strain showed a significant reduction of around 30% in the higher oligomeric crosslinked peptidoglycan (Fig. 7). But perhaps most importantly, the peptidoglycan isolated from the *sgtB/ltaS* double mutant S4-*sgtB* suppressor strain showed an increase in peptidoglycan crosslinking as compared to the original *ltaS* mutant strain (Fig. 7). We speculate that this increase in peptidoglycan cross-linking could potentially strengthen the cell wall to better sustain the high internal turgor pressure and be at least in part responsible for the observed growth improvement.

Bacterial two-hybrid studies have indicated that the *S. aureus* SgtB protein interacts with SgtA, PBP1, PBP2 and PBP2A (15). Therefore, SgtB and the main glycosyltransferase PBP2 might compete for substrate during peptidoglycan biosynthesis and inactivation of the former might increase the substrate availability and activity of the latter. This might in turn aid in the delivery of substrate to the transpeptidase domain of the bifunctional PBP2 enzyme resulting in increased crosslinking. Alternatively, SgtB could also affect the function of PBP4, which has been shown to be responsible for hyper-crosslinking of the staphylococcal peptidoglycan (53). Clearly, more experiments are needed to clarify the complex nature of peptidoglycan biosynthesis in *S. aureus* and changes observed in the absence of LTA.

Mutations in MazE could potentially also be tied in with changes in the cell wall structure of *S. aureus*. The MazEF type II toxin-anitoxin system is part of the *sigB* regulon and has been shown to be required for full activity of the alternative sigma factor SigB (37). It has also been reported that overexpression of σ^B^ causes cell wall thickening in *S. aureus* and increased resistance to cell wall-targeting antibiotics (38). Therefore, inactivation of MazE, as observed in one of our suppressor strains, could potentially also affect cell wall homeostasis via its effect on SigB.

In summary, our results suggest that in the absence of LTA, peptidoglycan in *S. aureus* becomes less cross-linked, which might weaken the cell wall and become less able to sustain the high internal turgor pressure, ultimately leading to cell lysis. The suppressor mutations obtained in *sgtB* (and perhaps also in some of the other genes observed in our screen) help the cell survive this detrimental effect by altering and strengthening the cell wall, likely allowing the cell wall to again better withstand the high internal turgor pressure.

## Materials and Methods

### Bacterial strains and culture conditions

All bacterial strains used in this study are listed in Table 3. *Escherichia coli* strains were cultured in Luria Bertani (LB) medium and *Staphylococcus aureus* strains in tryptic soya broth (TSB) at 37°C with aeration, unless otherwise stated. When appropriate the growth medium was supplemented with antibiotics and inducers as follow: for *E. coli* cultures: ampicillin (Amp) 100 µg/ml, chloramphenicol (Cam) 10 µg/ml, and kanamycin (Kan) 30 µg/ml; for *S. aureus* cultures: chloramphenicol (Cam) 10 µg/ml for plasmid selection and 7.5 µg/ml for chromosomally integrated plasmid selection, erythromycin (Erm) 10 µg/ml and kanamycin (Kan) 90 µg/ml. The inducer anhydrotetracycline (Atet) was used at a concentration of 100 or 200 ng/ml in agar plates and 50 ng/ml in broth.

**Table 3:**
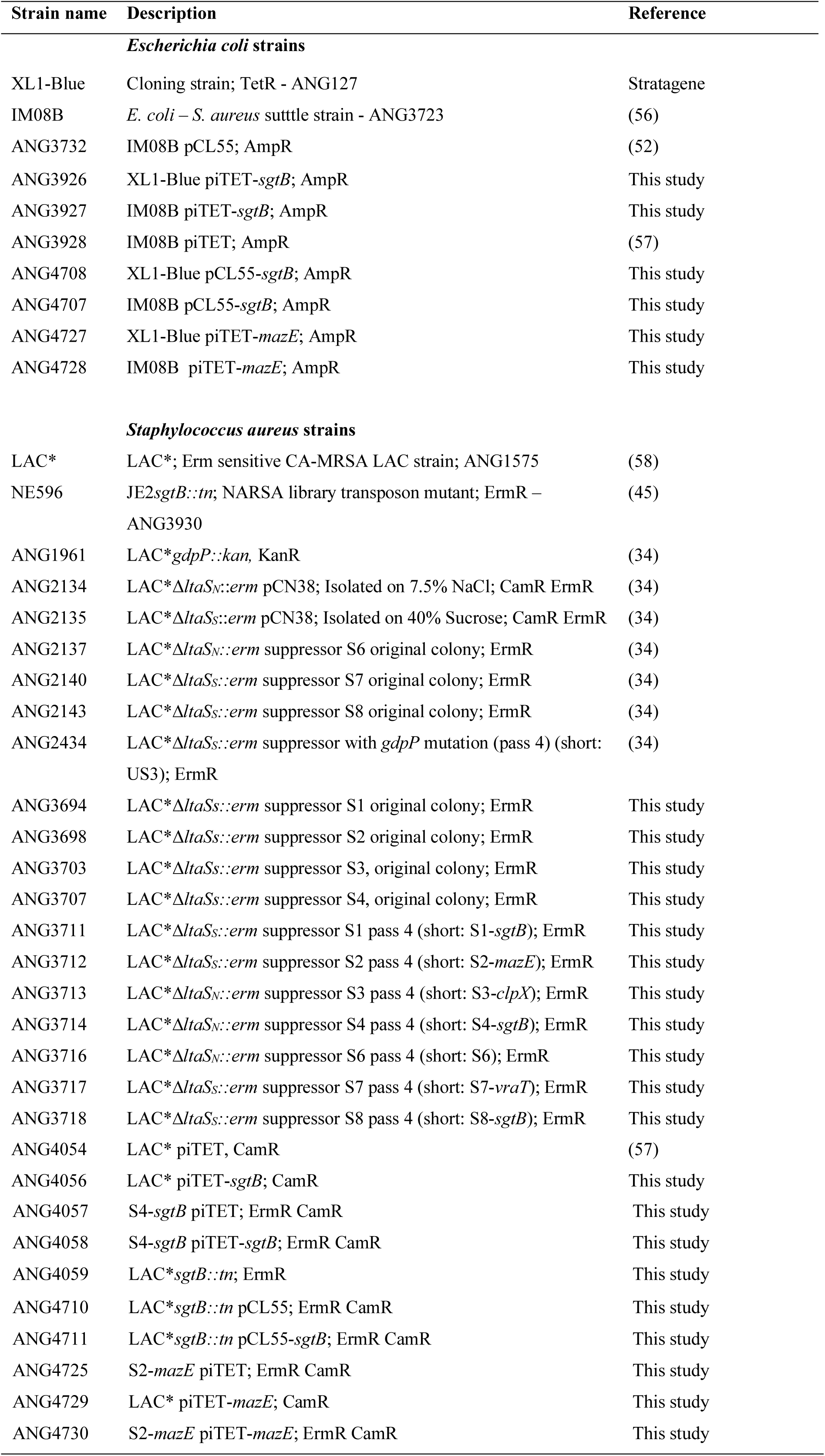
Bacterial strains used in this study

**Table 4:**
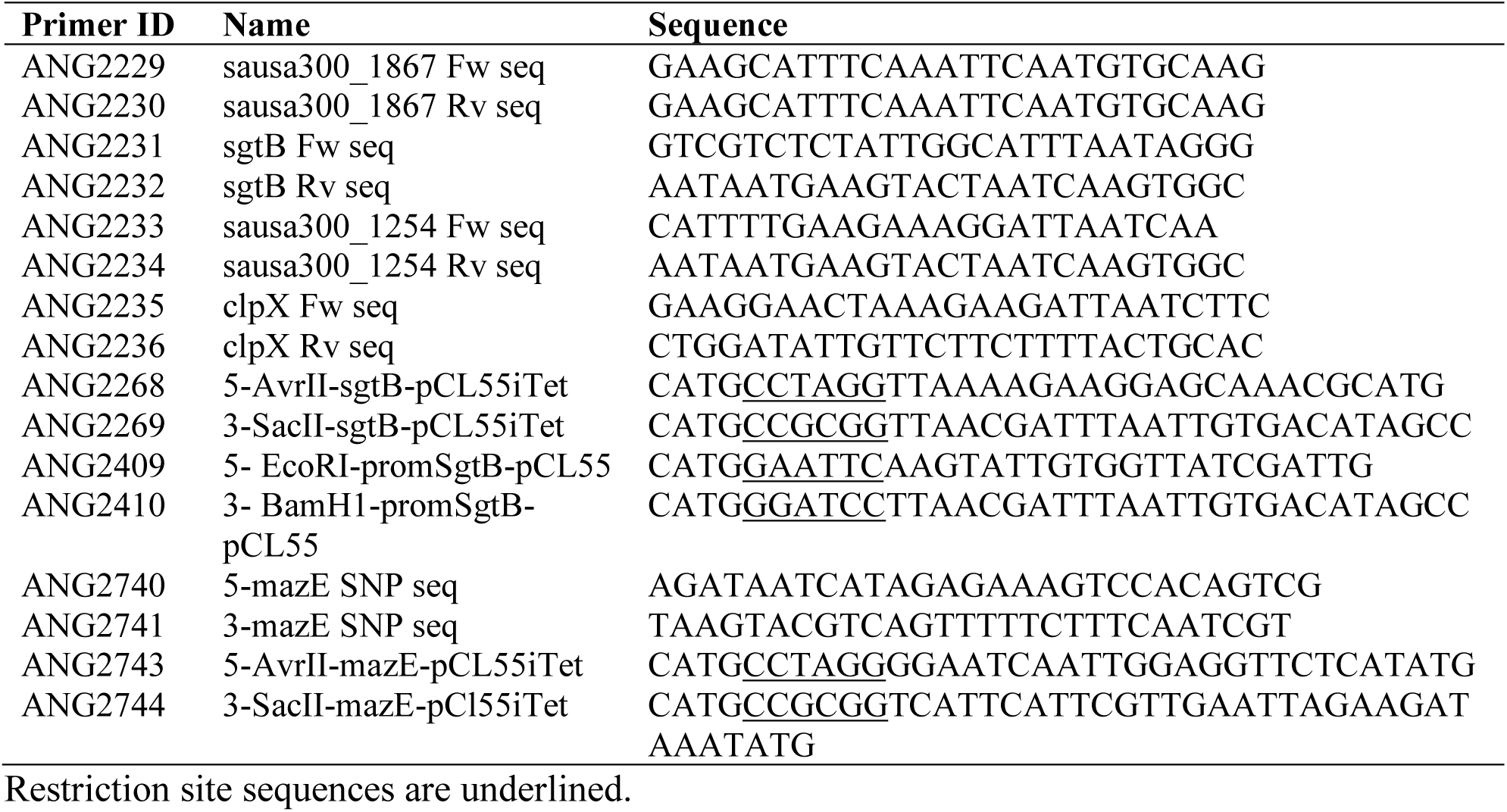
Primers used in this study

### Plasmid and strain construction

Strains and primers used in this study are listed in Tables 3 and 4, respectively. Strain LAC\**sgtB∷tn* was generated by transduction using phage Φ85 and transducing the *sgtB* region containing a transposon insertion in *sgtB* from the Nebraska transposon mutant library strain NE596 (45) into strain LAC*. Plasmids piTET-*sgtB* and piTET-*mazE* for anhydrotetracycline (Atet) inducible expression of *sgtB* and *mazE* in *S. aureus* were generated by amplifying *sgtB* or *mazE* from LAC* chromosomal DNA using primers pair ANG2268/ANG2269 or ANG2743/ANG2744, respectively. The PCR products and plasmid piTET were digested with AvrII and SacII, ligated and then transformed into *E. coli* XL1-Blue yielding strain XL1-Blue piTET-*sgtB* (ANG3926) and XL1-Blue piTET-*mazE* (ANG4727). The plasmids were shuttled through *E. coli* strain IM08B (ANG3927 and ANG4728) and subsequently electroporated into LAC* yielding strains LAC* piTET-*sgtB* (ANG4056) and LAC* piTET-*mazE* (ANG4729) or into the respective suppressor strains yielding strains S4-*sgtB* piTET-*sgtB* (ANG4058) and S2-*mazE* piTET-*mazE* (ANG4730). The plasmid pCL55-*sgtB* for complementation and expression of *sgtB* from its native promoter was generated by amplifying the *sgtB* gene including its native promoter region from LAC* chromosomal DNA using primers ANG2270 and ANG2271. The PCR product and plasmid pCL55 were digested with EcoRI and BamHI, ligated and then transformed into *E. coli* XL1-Blue yielding strain XL1-Blue pCL55-*sgtB* (ANG4708), shuttled through IM08B (ANG3923) and subsequently electroporated into the *sgtB* mutant strain yielding strain LAC\**sgtB∷tn* pCL55-*sgtB* (ANG4711). WT LAC* or mutant *S. aureus* strains with the empty vectors piTET or pCL55 (see Table 3) were used as control strains in several experiments. The sequences of all plasmid inserts were verified by automated fluorescence sequencing at GATC Biotechnology.

### Bacterial growth curves

*S. aureus* LAC* and the indicated *ltaS* suppressor strains were grown overnight in TSB medium containing the relevant antibiotic. The original *ltaS* mutant strains were grown in TSB containing either 7.5% NaCl or 40% sucrose. Overnight cultures were washed three times in TSB and diluted to a starting OD_600_ of 0.05. Cultures were incubated at 37°C with aeration and OD_600_ values determined at 2 h intervals. WT LAC* and the *sgtB* mutant strain LAC*\*sgtB∷tn* were grown overnight in TSB or TSB supplemented with 10 µg/ml Erm. The next day, bacteria were washed three times in TSB and diluted to a starting OD_600_ of 0.05 and 200 μl placed into a 96-well microtiter plate. Growth was monitored for 10h using a SPECTROSTARnano plate reader (BMG Labtech). All growth curves were performed in triplicates with average values and standard deviations plotted.

### Fluorescence microscopy analysis

Cells from 1 ml overnight culture were collected by centrifugation, washed 3 times in PBS pH 7.4 and 150 µl of these cell suspensions applied to polylysine (0.1%, w/v) treated coverslips. Bacteria-coated coverslips were incubated with 100 µl of a 1 µg/ml BODIPY-vancomycin (Molecular Probes) solution in H_2_O for 20 min, washed and mounted on glass slides containing 20 µl Vectashield (Vector Laboratories). Slides were viewed under a Zeiss Axio Imager A2, using a GFP filter set and images were captured with an AxioCam MRc Rev.3 camera and analyzed using the Zen Pro 2012 SP2 software. The experiment was performed in triplicate.

### Minimum Inhibitory Concentrations (MIC) determination

Overnight cultures of WT, the *sgtB* mutant and complementation stain as well as the indicated suppressor strains were grown over night in TSB medium. The next day, cultures were adjusted to an OD_600_ of 0.05 in TSB and 100 μl of these suspensions were incubated in 96-well plates with 2-fold dilutions of various antimicrobials at the following starting concentrations: oxacillin 500 µg/ml or 1 µg/ml as appropriate, daptomycin 32 µg/ml, lysostaphin 2 µg/ml, vancomycin 32 µg/ml and nisin 64 µg/ml. Oxacillin and daptomycin containing wells were supplemented with 2% (w/v) NaCl and 0.23 mM CaCl_2_, respectively. Plates were incubated at 37°C for 24 hours with shaking at 500 rpm. MICs were determined as the concentration of antibiotic at which growth was inhibited by >90% compared to growth without the antibiotic.

### LTA detection by western blot

LTA extraction and detection by western blot was performed as previously described (54). Briefly, samples were prepared from 1 ml overnight cultures normalized based on OD_600_ readings, that is cells from 1 ml culture with an OD_600_ of 6 were suspended in 90 µl 2 × SDS protein sample buffer. 10 µl of these samples were separated on 15% SDS-polyacrylamide gels and the material subsequently transferred to PVDF membranes. LTA was detected using the monoclonal polyglycerolphosphate-specific LTA antibody (Hycult Biotechnology) at a 1:4,000 dilution and the HRP-conjugated anti-mouse IgG antibody (Cell Signaling Technologies, USA) at a 1:10,000 dilution. The blots were developed by enhanced chemiluminescence using ClarityTM Western ECL Blotting Substrate (Bio-Rad) and imaged using the ChemiDoc Touch Imaging System (Bio-Rad). Western blots were performed in triplicates and a representative result is shown.

### Detection of WTA by Alcian blue silver staining

Flasks with 60 ml TSB medium supplemented with the appropriate antibiotics were inoculated with single colonies of WT LAC* and different suppressor strains and the cultures incubated overnight (18 h) with shaking at 37°C. The original *ltaS* mutant was grown in TSB supplemented with 40% sucrose and incubated 6-8 hours longer. Cells from an OD_600_ equivalent of 120 were harvested by centrifuging and the bacterial pellet stored at −20°C for further processing. Wall teichoic acid extraction and detection by Alcian blue silver staining following SDS PAGE analysis was performed as described in Covas *et al.* (55). Briefly, cells were washed in 20 ml buffer 1 (50 mM 2-(N-morpholino)ethanesulfonic acid (MES), pH 6.5), resuspended in 20 ml buffer 2 (buffer 1 with 4% (w/v) SDS) and boiled for 60 min. Next, the cells were washed with 20 ml buffer 2 and after transfer to 2.0 ml reaction tubes washed once more with 1.6 ml buffer 2, 1.6 ml buffer 3 (buffer 1 with 2% (w/v) NaCl) and finally with 1.6 ml buffer 1. The samples were suspended in 1.6 ml buffer 4 (20 mM Tris–HCl, pH 8.0, 0.5 % (w/v) SDS) and incubated for 4 h at 50°C with shaking following the addition of 2 µl of Proteinase K solution (20 mg /ml) from *Tritirachium album*. Next, the cells were collected by centrifugation, washed once with 1.6 ml buffer 3 and three times with 1.6 ml of H_2_O. To release the WTA, the pellets were suspended in 1 ml of 0.1 mM NaOH and the samples incubated for 16 h at 25°C. Next, the samples were centrifuged and 30 µl of the supernatant containing WTA separated on native Tris/Tricine polyacrylamide gels and the WTA visualized by Alcian blue silver staining (55).

### Whole genome sequencing

WT LAC*, LAC*Δ*ltaS*_*N*_, LAC*Δ*ltaS*_*S*_, and the indicated suppressor strains were cultured overnight at 37°C, cells harvested and genomic DNA extracted. Genome sequencing was performed by MicrobesNG (University of Birmingham) using an Illumina platform and a 250 bp paired end read kit. Sequence analysis was performed using the CLC Genomics work bench software package. First the LAC* reads were aligned against the published USA300 FPR3757 genome sequence (RefSeq accession number NC_007793.1), assembled into a reference contig and the USA300 FPR3757 annotation transferred onto the LAC* sequence. Next, the Illumina reads for the original *ltaS* mutant strains LAC*Δ*ltaS*_*S*_ and LAC*Δ*ltaS*_*N*_ as well as the seven different suppressor strains were mapped onto the assembled LAC* sequence and high frequency (> 65%) and good quality base changes identified using the CLC Genomics Workbench software package. Genomic alternations found in the suppressor strains but not present in the original *ltaS* mutant strain are summarized in Table 1. The Illumina short read for the WT LAC* has been previously published (35) and has been deposited in the European Nucleotide Archive under the study accession number PRJEB14759. The Illumina reads for the *ltaS* mutants and *ltaS* suppressor strains were deposited in the European Nucleotide Archive under the study accession number PRJEB14723.

### Peptidoglycan isolation and analysis

Overnight cultures of *S. aureus* strains LAC*, S4-*sgtB* and LAC\**sgtB∷tn* were prepared in TSB or TSB supplemented with 40% sucrose and of the original *ltaS* mutant strain LAC*Δ*ltaS*_*N*_ (ANG2134) in TSB supplemented with 40% sucrose. The next day, cells were back-diluted in 2 litres of the same growth medium to an OD_600_ of 0.05. The cultures were grown at 37°C until they reached an OD_600_ of approximately 1.5, cooled on ice and cells were collected by centrifugation. Peptidoglycan was purified and digested with mutanolysin as previously described (34, 48). HPLC analysis of the digested peptidoglycan material was performed as described previously (48) and the muropeptide profiles were determined for each strain and growth condition in triplicates. For the quantification of monomeric, dimeric, trimeric and higher oligomeric peptidoglycan material, the peaks were integrated. The total peak area for each muropetide profile was determined and set to 100% and the % monomeric, dimeric, trimeric, and higher oligomeric peaks calculated and the average values and standard deviation from the three profiles determined and plotted.

### c-di-AMP quantification by competitive ELISA

Five ml TSB were inoculated with a single colony of *S. aureus* LAC* (WT), strain LAC* *gdpP∷kan,* US3 (LAC*Δ*ltaS*_*S*_*∷erm* suppressor with mutation in *gdpP*) as well as the different LTA-negative suppressor and the tubes were incubated for 8 hours at 37°C. Next, the cultures were back-diluted to an OD_600_ of 0.05 in 10 ml TSB and grown for 15 hours at 37°C. Bacterial cells from these cultures were collected by centrifugation, cell lysates prepared and the cellular c-di-AMP levels determined using a previously described competitive ELISA method (35, 36). However, in place of determining c-di-AMP levels based on a standard curve the A_490_ values were directly compared. To this end, the A_490_ reading obtained for samples derived for the WT strain was set to 100% and the % values calculated for the samples derived from the other strains. Thee independent experiments were performed with three technical replicates and the average values and standard deviation of the % A_490_ values determined for each strain as compared to the WT strain. Of note, since this is a competitive ELISA, and a decrease in A_490_ readings represents an increase in cellular c-di-AMP levels.

## Acknowledgments

This research was supported by the European Research Council grant 260371 and the Wellcome Trust grant 100289 to AG, the UK Medical Research Council (MRC) within the AMR Cross-council initiative Collaborative Grant MR/N002679/1 to WV and the German Research Foundation [Deutsche Forschungsgemeinschaft (DFG)] grant SCHU 3159/1-1 to CFS. The Illumina sequencing was performed by MicrobesNG, which was supported by the BBSRC grant BB/L024209/1.

## REFERENCES

1. Kluytmans J, Van Belkum A, Verbrugh H. 1997. Nasal carriage of *Staphylococcus aureus*: epidemiology, underlying mechanisms, and associated risks. Clinical microbiology reviews 10:505–520.

2. Baek KT, Bowman L, Millership C, Dupont Sogaard M, Kaever V, Siljamaki P, Savijoki K, Varmanen P, Nyman TA, Gründling A, Frees D. 2016. The Cell Wall Polymer Lipoteichoic Acid Becomes Nonessential in *Staphylococcus aureus* Cells Lacking the ClpX Chaperone. MBio 7.

3. Francis JS, Doherty MC, Lopatin U, Johnston CP, Sinha G, Ross T, Cai M, Hansel NN, Perl T, Ticehurst JR. 2005. Severe community-onset pneumonia in healthy adults caused by methicillin-resistant *Staphylococcus aureus* carrying the Panton-Valentine leukocidin genes. Clinical Infectious Diseases 40:100–107.

4. S. Hiramatsu K, Katayama Y, Matsuo M, Sasaki T, Morimoto Y, Sekiguchi A, Baba 2014. Multi-drug-resistant *Staphylococcus aureus* and future chemotherapy. Journal of Infection and Chemotherapy 20:593–601.

5. Horn J, Stelzner K, Rudel T, Fraunholz M. 2017. Inside job: *Staphylococcus aureus* host-pathogen interactions. International Journal of Medical Microbiology.

6. Foster TJ, Geoghegan JA, Ganesh VK, Höök M. 2014. Adhesion, invasion and evasion: the many functions of the surface proteins of *Staphylococcus aureus*. Nat Rev Microbiol 12:49–62.

7. Silhavy TJ, Kahne D, Walker S. 2010. The bacterial cell envelope. Cold Spring Harbor perspectives in biology 2:a000414.

8. Rajagopal M, Walker S. 2015. Envelope structures of gram-positive bacteria, p 1–44, Protein and Sugar Export and Assembly in Gram-positive Bacteria. Springer.

9. Vollmer W, Blanot D, De Pedro MA. 2008. Peptidoglycan structure and architecture. FEMS microbiology reviews 32:149–167.

10. Archibald A, Hancock I, Harwood C. 1993. Cell wall structure, synthesis, and turnover, p 381–410, Bacillus subtilis and other Gram-positive bacteria. American Society of Microbiology.

11. Holtje JV. 1998. Growth of the stress-bearing and shape-maintaining murein sacculus of *Escherichia coli*. Microbiol Mol Biol Rev 62:181–203.

12. Reed P, Atilano ML, Alves R, Hoiczyk E, Sher X, Reichmann NT, Pereira PM, Roemer T, Filipe SR, Pereira-Leal JB, Ligoxygakis P, Pinho MG. 2015. *Staphylococcus aureus* Survives with a Minimal Peptidoglycan Synthesis Machine but Sacrifices Virulence and Antibiotic Resistance. PLoS Pathog 11:e1004891.

13. Hartman BJ, Tomasz A. 1984. Low-affinity penicillin-binding protein associated with beta-lactam resistance in *Staphylococcus aureus*. J Bacteriol 158:513–516.

14. Pinho MG, de Lencastre H, Tomasz A. 2001. An acquired and a native penicillin-binding protein cooperate in building the cell wall of drug-resistant staphylococci. Proc Natl Acad Sci U S A 98:10886–10891.

15. Reed P, Veiga H, Jorge AM, Terrak M, Pinho MG. 2011. Monofunctional transglycosylases are not essential for *Staphylococcus aureus* cell wall synthesis. J Bacteriol 193:2549–2556.

16. Wang QM, Peery RB, Johnson RB, Alborn WE, Yeh WK, Skatrud PL. 2001. Identification and characterization of a monofunctional glycosyltransferase from *Staphylococcus aureus*. J Bacteriol 183:4779–4785.

17. Weidenmaier C, Peschel A. 2008. Teichoic acids and related cell-wall glycopolymers in Gram-positive physiology and host interactions. Nature Reviews Microbiology 6:276.

18. Kawai Y, Marles-Wright J, Cleverley RM, Emmins R, Ishikawa S, Kuwano M, Heinz N, Bui NK, Hoyland CN, Ogasawara N, Lewis RJ, Vollmer W, Daniel RA, Errington J. 2011. A widespread family of bacterial cell wall assembly proteins. Embo j 30:4931–4941.

19. Kern T, Giffard M, Hediger S, Amoroso A, Giustini C, Bui NK, Joris B, Bougault C, Vollmer W, Simorre JP. 2010. Dynamics characterization of fully hydrated bacterial cell walls by solid-state NMR: evidence for cooperative binding of metal ions. J Am Chem Soc 132:10911–10919.

20. Neuhaus FC, Baddiley J. 2003. A continuum of anionic charge: structures and functions of D-alanyl-teichoic acids in gram-positive bacteria. Microbiology and Molecular Biology Reviews 67:686–723.

21. Brown S, Santa Maria JP, Jr., Walker S. 2013. Wall teichoic acids of gram-positive bacteria. Annu Rev Microbiol 67:313–336.

22. Percy MG, Gründling A. 2014. Lipoteichoic acid synthesis and function in gram-positive bacteria. Annu Rev Microbiol 68:81–100.

23. Percy MG, Karinou E, Webb AJ, Gründling A. 2016. Identification of a Lipoteichoic Acid Glycosyltransferase Enzyme Reveals that GW-Domain-Containing Proteins Can Be Retained in the Cell Wall of Listeria monocytogenes in the Absence of Lipoteichoic Acid or Its Modifications. J Bacteriol 198:2029–2042.

24. Santa Maria JP, Jr., Sadaka A, Moussa SH, Brown S, Zhang YJ, Rubin EJ, Gilmore MS, Walker S. 2014. Compound-gene interaction mapping reveals distinct roles for *Staphylococcus aureus* teichoic acids. Proc Natl Acad Sci U S A 111:12510–12515.

25. Peschel A, Otto M, Jack RW, Kalbacher H, Jung G, Götz F. 1999. Inactivation of the dlt Operon in *Staphylococcus aureus* Confers Sensitivity to Defensins, Protegrins, and Other Antimicrobial Peptides. Journal of Biological Chemistry 274:8405–8410.

26. Kho K, Meredith TC. 2018. Salt-Induced Stress Stimulates a Lipoteichoic Acid-Specific Three Component Glycosylation System in *Staphylococcus aureus*. J Bacteriol doi:10.1128/JB.00017-18.

27. Reichmann NT, Gründling A. 2011. Location, synthesis and function of glycolipids and polyglycerolphosphate lipoteichoic acid in Gram-positive bacteria of the phylum Firmicutes. FEMS Microbiol Lett 319:97–105.

28. Lu D, Wörmann ME, Zhang X, Schneewind O, Gründling A, Freemont PS. 2009. Structure-based mechanism of lipoteichoic acid synthesis by *Staphylococcus aureus* LtaS. Proceedings of the National Academy of Sciences 106:1584–1589.

29. Gründling A, Schneewind O. 2007. Synthesis of glycerol phosphate lipoteichoic acid in *Staphylococcus aureus*. Proc Natl Acad Sci U S A 104:8478–8483.

30. Karatsa-Dodgson M, Wormann ME, Gründling A. 2010. In vitro analysis of the *Staphylococcus aureus* lipoteichoic acid synthase enzyme using fluorescently labeled lipids. J Bacteriol 192:5341–5349.

31. Oku Y, Kurokawa K, Matsuo M, Yamada S, Lee BL, Sekimizu K. 2009. Pleiotropic roles of polyglycerolphosphate synthase of lipoteichoic acid in growth of *Staphylococcus aureus* cells. J Bacteriol 191:141–151.

32. Schirner K, Marles-Wright J, Lewis RJ, Errington J. 2009. Distinct and essential morphogenic functions for wall-and lipo-teichoic acids in *Bacillus subtilis*. The EMBO journal 28:830–842.

33. Fedtke I, Mader D, Kohler T, Moll H, Nicholson G, Biswas R, Henseler K, Götz F, Zähringer U, Peschel A. 2007. A *Staphylococcus aureus ypfP* mutant with strongly reduced lipoteichoic acid (LTA) content: LTA governs bacterial surface properties and autolysin activity. Molecular microbiology 65:1078–1091.

34. Corrigan RM, Abbott JC, Burhenne H, Kaever V, Gründling A. 2011. c-di-AMP is a new second messenger in *Staphylococcus aureus* with a role in controlling cell size and envelope stress. PLoS Pathog 7:e1002217.

35. Bowman L, Zeden MS, Schuster CF, Kaever V, Gründling A. 2016. New Insights into the Cyclic Di-adenosine Monophosphate (c-di-AMP) Degradation Pathway and the Requirement of the Cyclic Dinucleotide for Acid Stress Resistance in *Staphylococcus aureus*. J Biol Chem 291:26970–26986.

36. Underwood AJ, Zhang Y, Metzger DW, Bai G. 2014. Detection of cyclic di-AMP using a competitive ELISA with a unique pneumococcal cyclic di-AMP binding protein. Journal of microbiological methods 107:58–62.

37. Donegan NP, Cheung AL. 2009. Regulation of the mazEF toxin-antitoxin module in *Staphylococcus aureus* and its impact on sigB expression. Journal of bacteriology 191:2795–2805.

38. Morikawa K, Maruyama A, Inose Y, Higashide M, Hayashi H, Ohta T. 2001. Overexpression of sigma factor, sigma(B), urges *Staphylococcus aureus* to thicken the cell wall and to resist beta-lactams. Biochem Biophys Res Commun 288:385–389.

39. Guldimann C, Boor KJ, Wiedmann M, Guariglia-Oropeza V. 2016. Resilience in the face of uncertainty: sigma factor B fine-tunes gene expression to support homeostasis in gram-positive bacteria. Applied and environmental microbiology 82:4456–4469.

40. Kuroda M, Kuroda H, Oshima T, Takeuchi F, Mori H, Hiramatsu K. 2003. Two-component system VraSR positively modulates the regulation of cell-wall biosynthesis pathway in *Staphylococcus aureus*. Mol Microbiol 49:807–821.

41. Kuroda M, Kuwahara-Arai K, Hiramatsu K. 2000. Identification of the up-and down-regulated genes in vancomycin-resistant *Staphylococcus aureus* strains Mu3 and Mu50 by cDNA differential hybridization method. Biochem Biophys Res Commun 269:485–490.

42. Boyle-Vavra S, Yin S, Jo DS, Montgomery CP, Daum RS. 2013. VraT/YvqF is required for methicillin resistance and activation of the VraSR regulon in *Staphylococcus aureus*. Antimicrob Agents Chemother 57:83–95.

43. DeFrancesco AS, Masloboeva N, Syed AK, DeLoughery A, Bradshaw N, Li GW, Gilmore MS, Walker S, Losick R. 2017. Genome-wide screen for genes involved in eDNA release during biofilm formation by *Staphylococcus aureus*. Proc Natl Acad Sci U S A 114:E5969–E5978.

44. Vickery CR, Wood BM, Morris HG, Losick R, Walker S. 2018. Reconstitution of *Staphylococcus aureus* Lipoteichoic Acid Synthase Activity Identifies Congo Red as a Selective Inhibitor. J Am Chem Soc 140:876–879.

45. Fey PD, Endres JL, Yajjala VK, Widhelm TJ, Boissy RJ, Bose JL, Bayles KW. 2013. A genetic resource for rapid and comprehensive phenotype screening of nonessential *Staphylococcus aureus* genes. MBio 4:e00537–00512.

46. Terrak M, Nguyen-Distèche M. 2006. Kinetic characterization of the monofunctional glycosyltransferase from *Staphylococcus aureus*. Journal of bacteriology 188:2528–2532.

47. Rebets Y, Lupoli T, Qiao Y, Schirner K, Villet R, Hooper D, Kahne D, Walker S. 2014. Moenomycin resistance mutations in *Staphylococcus aureus* reduce peptidoglycan chain length and cause aberrant cell division. ACS Chem Biol 9:459–467.

48. De Jonge B, Chang Y-S, Gage D, Tomasz A. 1992. Peptidoglycan composition of a highly methicillin-resistant Staphylococcus aureus strain. The role of penicillin binding protein 2A. Journal of Biological Chemistry 267:11248–11254.

49. Reichmann NT, Picarra Cassona C, Monteiro JM, Bottomley AL, Corrigan RM, Foster SJ, Pinho MG, Gründling A. 2014. Differential localization of LTA synthesis proteins and their interaction with the cell division machinery in *Staphylococcus aureus*. Mol Microbiol 92:273–286.

50. Commichau FM, Gibhardt J, Halbedel S, Gundlach J, Stülke J. 2017. A Delicate Connection: c-di-AMP Affects Cell Integrity by Controlling Osmolyte Transport. Trends in microbiology 26:175–185.

51. Corrigan RM, Campeotto I, Jeganathan T, Roelofs KG, Lee VT, Gründling A. 2013. Systematic identification of conserved bacterial c-di-AMP receptor proteins. Proceedings of the National Academy of Sciences 110:9084–9089.

52. Schuster CF, Bellows LE, Tosi T, Campeotto I, Corrigan RM, Freemont P, Gründling A. 2016. The second messenger c-di-AMP inhibits the osmolyte uptake system OpuC in *Staphylococcus aureus*. Sci Signal 9:ra81.

53. Memmi G, Filipe SR, Pinho MG, Fu Z, Cheung A. 2008. *Staphylococcus aureus* PBP4 is essential for beta-lactam resistance in community-acquired methicillin-resistant strains. Antimicrob Agents Chemother 52:3955–3966.

54. Gründling A, Schneewind O. 2007. Genes required for glycolipid synthesis and lipoteichoic acid anchoring in *Staphylococcus aureus*. J Bacteriol 189:2521–2530.

55. Covas G, Vaz F, Henriques G, Pinho M, Filipe S. 2016. Analysis of Cell Wall Teichoic Acids in *Staphylococcus aureus*., p 201–213. *In* Hong H (ed), Bacterial Cell Wall Homeostasis, vol 1440. Humana Press, New York, NY.

56. Monk IR, Tree JJ, Howden BP, Stinear TP, Foster TJ. 2015. Complete Bypass of Restriction Systems for Major *Staphylococcus aureus* Lineages. MBio 6:e00308–00315.

57. Zeden MS, Schuster CF, Bowman L, Zhong Q, Williams HD, Gründling A. 2018. Cyclic-di-adenosine monophosphate (c-di-AMP) is required for osmotic regulation in *Staphylococcus aureus* but dispensable for viability in anaerobic conditions. J Biol Chem doi:10.1074/jbc.M117.818716.

58. Boles BR, Thoendel M, Roth AJ, Horswill AR. 2010. Identification of genes involved in polysaccharide-independent *Staphylococcus aureus* biofilm formation. PLoS One 5:e10146.

